# Dual Role of Ninjurin-1 in Myeloid Cell Adhesion and Inflammation in Relapse-Remitting EAE

**DOI:** 10.64898/2026.01.16.699993

**Authors:** Coleen Thompson, Alex Annett, Anastasia Alkhimovitch, Kashish Singh Parihar, Igal Ifergan

**Affiliations:** Department of Molecular and Cellular Biology, College of Medicine, University of Cincinnati, Cincinnati, OH USA; Division of Immunobiology, Cincinnati Children’s Hospital Medical Center, University of Cincinnati College of Medicine, Cincinnati, OH USA; Neuroscience Graduate Program, College of Medicine, University of Cincinnati, Cincinnati, OH USA

**Keywords:** Ninjurin-1, EAE, myeloid cells, multiple sclerosis, CNS inflammation

## Abstract

Nerve Injury-Induced Protein 1 (Ninjurin-1) is an adhesion molecule implicated in inflammation and tissue injury, yet its role in neuroinflammatory diseases such as multiple sclerosis (MS) remains poorly defined. Here, we identify Ninjurin-1 as a key mediator of immune activation and CNS infiltration in relapsing-remitting experimental autoimmune encephalomyelitis (RR-EAE), a model of relapsing-remitting MS (RRMS). Using flow cytometry, gene-expression profiling, and *in vivo* peptide blockade, we show that Ninjurin-1 is markedly upregulated on CNS-infiltrating myeloid cells during disease progression. Ninjurin-1⁺ myeloid cells display a dual function, as both an adhesion molecule and a marker of inflammatory activation, characterized by increased antigen presentation, cytokine production, and transcriptional enrichment for genes regulating adhesion, migration, and innate immune signaling. Importantly, therapeutic blockade of Ninjurin-1 significantly reduced clinical severity, CNS immune infiltration, and demyelination in RR-EAE. These findings uncover a previously unrecognized role for Ninjurin-1 in myeloid-driven neuroinflammation and highlight its potential as a therapeutic target for relapsing-remitting MS.

## 1. Introduction

Multiple sclerosis (MS) is a chronic autoimmune disorder affecting over 2.8 million people worldwide, leading to progressive neurological disability and reduced quality of life [1, 2]. The disease is driven by the infiltration of peripheral immune cells into the central nervous system (CNS), where they attack the myelin sheath surrounding neurons, causing demyelination, inflammation, and neurodegeneration [3]. Although the precise etiology of MS remains unclear, a prevailing hypothesis proposes that antigen-presenting cells (APCs) present myelin-mimicking antigens to autoreactive T lymphocytes, thereby initiating immune-mediated demyelination and lesion formation within the CNS [4, 5].

Experimental autoimmune encephalomyelitis (EAE), a well-established preclinical animal model that recapitulates many clinical and pathological features of MS, including limb paralysis, CNS lesions, and immune cell infiltration [6, 7]. EAE studies have identified key roles for TH1 and TH17 CD4⁺ T cells in driving neuroinflammation and demyelination [8], providing a valuable model to investigate disease mechanisms and test potential therapies.

Immune cell migration into the CNS is orchestrated by a complex interplay of adhesion molecules, chemokines, and selectins expressed by blood–brain barrier endothelial cells (BBB-ECs) and CD45⁺ immune cells [9]. Among these, vascular cell adhesion molecule-1 (VCAM-1) and intercellular adhesion molecule-1 (ICAM-1) are well-established mediators of leukocyte trafficking in MS and EAE [10]. However, the molecular mechanisms governing immune cell infiltration into the CNS remain incompletely understood, suggesting that additional adhesion pathways contribute to neuroinflammatory progression.

One such candidate is Nerve Injury–Induced Protein-1 (Ninjurin-1), a 22-kDa homophilic adhesion molecule originally identified for its role in axonal regeneration following nerve injury [11]. Subsequent work, including a proteomic screen of human BBB-ECs by Ifergan et al. (2013), identified Ninjurin-1 as a molecule expressed at the neurovascular interface and upregulated under inflammatory conditions [12]. Importantly, Ninjurin-1 is also expressed by immune cells, positioning it as a bidirectional mediator of leukocyte-endothelial interactions.

More recently, Ninjurin-1 has emerged as a regulator of immune cell adhesion and inflammation across pathological contexts, including stroke [13], pulmonary fibrosis [14], liver ischemia-reperfusion injury [15], and MS [12]. Beyond its adhesive function, Ninjurin-1 promotes leukocyte adhesion, plasma membrane rupture (PMR), and the release of damage-associated molecular patterns (DAMPs) [16], highlighting its potential role in amplifying inflammatory cascades. These findings position Ninjurin-1 as a unique molecule bridging structural adhesion and innate immune activation.

In the context of MS and EAE, Ninjurin-1 is expressed in both acute and chronic MS lesions [17] and is upregulated on myeloid and endothelial cells during chronic EAE [12]. Our previous work demonstrated that Ninjurin-1 expression peaks at the height of disease in a chronic EAE model induced by MOG_35-55_, and that its blockade alleviates clinical symptoms, reduces CNS inflammation, and limits immune cell infiltration [12]. Despite these insights, no studies have explored the role of Ninjurin-1 in relapse-remitting EAE (RR-EAE) or its relevance to relapse-remitting MS (RRMS). RRMS is the most prevalent MS subtype, representing approximately 85% of all cases. It is characterized by episodic neuroinflammatory attacks followed by partial recovery, driven by waves of peripheral immune cell infiltration into the CNS [18, 19].

In this study, we investigate the expression and function of Ninjurin-1 in RR-EAE. We show that Ninjurin-1 is strongly upregulated in inflammatory environments, particularly on myeloid cells and BBB-ECs, where it functions both as an adhesion molecule and as a marker of inflammation. Furthermore, blockade of Ninjurin-1 homophilic binding attenuates disease severity and reduces immune cell infiltration. Together, these findings identify Ninjurin-1 as a dual-function mediator of immune activation and trafficking in RR-EAE and suggest that targeting Ninjurin-1 may represent a promising therapeutic strategy for RRMS.

## 2. Materials and Methods

### 2.1 Mice

Female SJL/J (cat# 000686) and C57BL/6 (cat# 000664) mice from 6-9 weeks old were purchased from The Jackson Laboratory (Bar Harbor, ME). 2D2 mice were bred in-house (originally from The Jackson Laboratory, cat# 006912) and used in experiments once they reached 12-16 weeks old. All CD4+ T cells from 2D2 mice possess a T cell receptor specific (TCR) for MOG_35-55_. All mice were maintained under specific pathogen-free conditions in the University of Cincinnati’s Laboratory Animal Medical Services (LAMS) facility and handled in accordance with AAALAC and IACUC-approved institutional guidelines.

### 2.2 Reagents

Peptides corresponding to the adhesion motif of Ninjurin-1 (amino acids 26-37; sequence PPRWGLRNRPIN) were synthesized and used as a blocking peptide (anti-Ninj_26-37_). A scrambled control peptide (WRGNPGIRWAPH) was also generated for use as a control. Peptides were synthesized by Genemed Biotechnologies Inc. (Torrance, CA).

### 2.3 Active EAE

RR-EAE was induced in 8-10-week-old female SJL/J mice as previously described [20]. Mice received three subcutaneous flank injections (total 100 µL) of PLP_139-151_ peptide (200 µg/mouse; Genemed Biotechnologies Inc.) emulsified in Complete Freund’s Adjuvant (BD Biosciences) containing 200 µg of *Mycobacterium tuberculosis* H37Ra. At peak of disease, mice were treated intraperitoneally three times per week with 200 µg of either scrambled peptide or Ninj_26–37_ blocking peptide (1 mg/mL) until the study endpoint (n = 8/group). Clinical scores were recorded daily (0 = asymptomatic; 1 = limp tail; 2 = hind limb weakness; 3 = partial paralysis; 4 = complete paralysis; 5 = moribund). Data are presented as mean ± SEM daily score.

### 2.4 Isolation of splenocytes and CNS

Mice were incapacitated with CO_2_ and then perfused with PBS prior to tissue removal, following previously described methods [18]. Spleens were collected at pre-onset, onset, peak, remission, and relapse phases of RR-EAE. Tissue was mechanically dissociated through 100 µm cell strainers and red blood cells were lysed using 0.83% ammonium chloride. Cells were then plated in 96-well plates (0.2-2 x 10^6^ cells/well) in RPMI-1640 supplemented with 10% FBS, 2 mM L-glutamine, and 100 U/mL penicillin-streptomycin at 37 °C, 5% CO_2_.

CNS tissues (brain and spinal cord) were collected at the same time as the spleen and digested with 50 µg/mL DNase I and 500 µg/mL collagenase (Sigma). Mononuclear cells were isolated by 40% Percoll gradient or 20% BSA centrifugation. White blood cells were quantified using a Sysmex XP300 (Horst, IL).

### 2.5 Ex vivo cytokine stimulation

CD11b⁺ cells were enriched from naïve SJL/J splenocytes using MACS Magnetic separation columns (Miltenyi Biotec) and were plated at 0.5 x 10^6^ cells/well in 24 well plates. C57BL/6 brain microvascular endothelial cells (Cell Biologics, C57-6023) were cultured to 90% confluency and then plated at 30,000 cells/well in a 24 well plate. Cells were treated for 24 hours with the following cytokines: GM-CSF (200 ng/mL; R&D), IFN-γ (200 ng/mL; R&D), IL-1β (200 ng/mL; R&D), IL-4 (200 ng/mL; Peprotech), IL-6 (50 ng/mL; R&D), IL-10 (200 ng/mL; R&D), IL-17 (200 ng/mL; R&D), TGF-β (200 ng/mL; R&D), or TNF-α (50 ng/mL; R&D). Combination conditions included IFN-γ + TNF-α (referred to as TH1-like) and IL-17 + GM-CSF (referred to as TH17-like). Cytokine concentrations were based on manufacturer-reported ED_50_ values. After the 24-hour incubation (37°C, 5% CO_2_), Ninjurin-1 expression was analyzed by flow cytometry.

### 2.6 Flow cytometry staining and analysis

Fc receptors were blocked using anti-mouse CD16/32 (0.25 μg; Thermofisher). Cells were then stained in PBS with fixable LIVE/DEAD reagents (Life Technologies) for 20 minutes in PBS at room temperature in the dark to assess viability. Then cells were stained for surface markers for 20 minutes at 4°C using the specified antibodies (Ninjurin-1 from R&D Systems; all the others from BD Biosciences or Biolegend). When needed, cells were fixed and permeabilized (ThermoFisher) to stain for intracellular markers. To detect T cell cytokine expression, cells were activated for 4 hours with 1 mg/ml ionomycin (Iono) and 20 ng/ml phorbol 12-myristate 13-acetate 40 (PMA) in the presence of 2 mg/ml brefeldin A (BFA) (Sigma). To detect CD11b^+^ cytokine expression, cells were activated for 18 hours with 100 ng/ml lipopolysaccharide (LPS) from E. coli serotype 0111:B4 (Sigma) in the presence of 2 mg/ml BFA (Sigma) for the last 2 hours of co-culture. Cells were stained for surface markers and a fixation and permeabilization kit (ThermoFisher) was used. Cells were acquired using a BD FACSCanto II and analyzed using Flowjo version 10.1 software.

### 2.7 Immunostaining of mouse CNS

CNS tissues from control and RR-EAE mice were fixed in 10% formaldehyde overnight. Luxol Fast Blue (LFB) and hematoxylin and eosin (H&E) staining were performed by the Immunohistochemistry (IHC) Core at the University of Cincinnati. Images were acquired with an EVOS M5000 microscope (Invitrogen) at 10x magnification.

For quantification of cellular infiltration in H&E-stained sections, hematoxylin-positive nuclei were manually counted within standardized white matter regions using ImageJ. Three non-overlapping regions of interest (ROIs; 300 × 300 µm each) were analyzed per section, and counts were averaged to generate a single value per section. Values from multiple sections were then averaged to yield one representative value per mouse. Data are expressed as cells per mm².

### 2.8 Cell sorting

Splenocytes were removed from naïve C57BL/6 mice and enriched with CD11b magnetic beads from Miltenyi Biotec. Cells were then stained with Ninjurin-1, CD45, CD11b, CD3, B220, Ly6G, and Live Dead antibodies (**Fig. S4**) and sorted into two populations, CD45^+^CD11b^+^B220^-^CD3^-^Ly6G^-^Ninjurin-1^+^ (Ninjurin-1^+^) or CD45^+^CD11b^+^B220^-^CD3^-^Ly6G^-^Ninjurin-1^-^ (Ninjurin-1^-^) using a MA900 (Sony).

### 2.9 Quantitative PCR

After cell sorting, both populations obtained from cell sorting (Ninjurin-1^+^ and Ninjurin-1^-^) were lysed using Trizol (Ambion). Purity of the isolated RNA was determined by measuring the ratio of the optical density of the samples at 260/280nm using a Nanodrop spectrophotometer (Thermo Scientific). The OD_260_/OD_280_ ratio ranged from 1.7 to 2.1 for all samples. cDNAs were synthesized using the RT^2^ First Strand kit (Qiagen) according to the manufacturer’s instructions. The RT^2^ Profiler PCR Array Mouse Dendritic and Antigen Presenting Cell (PAMM-406Z) plates were purchased from SABiosciences, Qiagen. This array profiles the expression of 84 genes involved in antigen presentation and includes 5 controls for housekeeping genes, one control for genomic DNA, and three reverse transcription controls. PCRs were performed on a QuantStudio 3 (Thermofisher). The data were analyzed using the web-based software RT^2^ Profiler PCR Array data analysis tool (Qiagen). The C_T_ cut off was 35 and the data was normalized using Beta-2 microglobulin (B2m), Heat shock protein 90 alpha (cytosolic), and class B member 1 (HSP90ab1) housekeeping genes. Fold-changes for each gene were calculated as the difference in gene expression between the Ninjurin-1^-^ cells and Ninjurin-1^+^ cells. A positive value indicates gene up-regulation, and a negative value indicated gene down-regulation on the Ninjuirin-1^+^ cells. Only fold changes of 1.5 or greater were considered in our analysis.

### 2.8 2D2 Co-culture

Splenocytes were isolated from 2D2 mice, and CD4^+^ cells were magnetically separated via a negative CD4^+^ selection kit (Stemcell) and then labeled with carboxyfluorescein succinimidyl ester (CFSE) (Invitrogen). CFSE^+^CD4^+^ T cells (100,000/well) were then co-cultured with either Ninjurin-1^-^ or Ninjurin-1^+^ sorted cells (40,000/well) with 20 μg/ml MOG_35-55_ peptide (Genemed Biotechnologies Inc.) into a 96 well plate for 72 hours. After 72 hours, PMA/Iono/BFA was added for 4 hours to the culture prior to cell staining. The expression of CFSE, CD25, GM-CSF, IL-17, IFNγ, and TNF-α on T cells was analyzed via flow cytometry.

### 2.9 Statistical analysis

Statistical analyses were performed using GraphPad PRISM 10.0 (GraphPad software). Data are presented as the mean ± the standard error of the mean (SEM). EAE scores were analyzed by nonparametric Mann-Whitney test. All other analyses were performed by a paired t-test or a two-way Anova. Only *p* values < 0.05 were considered significant.

## 3. Results

### 3.1 Myeloid cells upregulate Ninjurin-1 expression in the CNS during RR-EAE

We sought to determine whether Ninjurin-1 played a role in RR-EAE, a mouse model of RRMS, as this had not been previously investigated. To assess the dynamics of Ninjurin-1 expression over the course of RR-EAE, we analyzed immune cells isolated from the CNS (brain and spinal cord) and spleen using flow cytometry at defined disease stages: pre-onset (days 7-9), onset (days 11-13), peak (days 14-16), remission (days 21-23), and relapse (days 28-32) (**Fig. 1A**). RR-EAE was induced in female SJL/J mice with PLP_139-151_. Our analysis focused on CD11b⁺ myeloid cells and CD3⁺ T cells, the principal immune cell populations infiltrating the CNS during RR-EAE [21–23]. The gating strategy can be found be found in (**Fig. S1)**.

**Figure 1.**
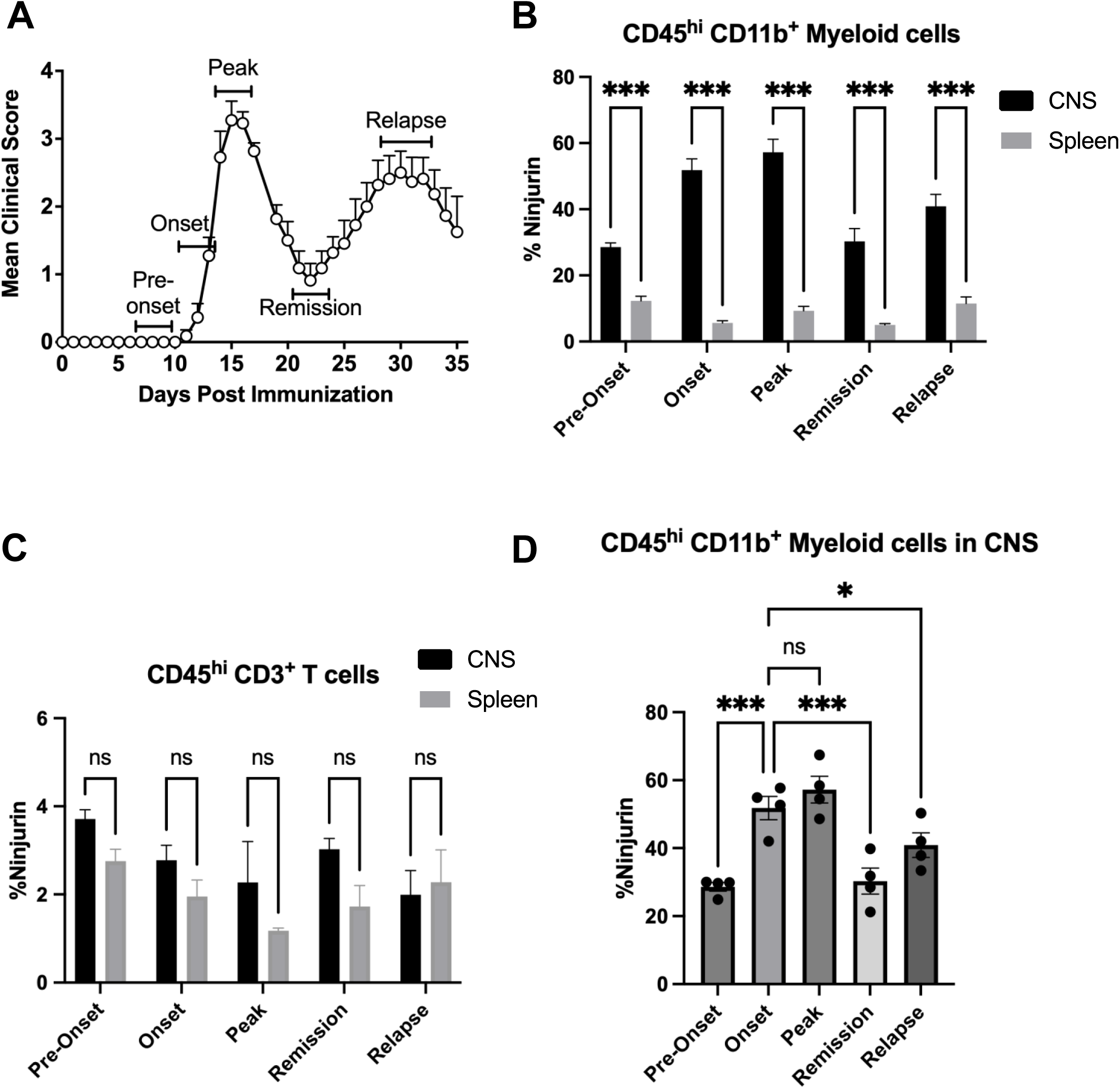
Ninjurin-1 is highly expressed on myeloid cells in the CNS throughout RR-EAE. RR-EAE was induced in female SJL/J mice using PLP_139-151_ emulsified in complete Freund’s adjuvant. Spleen and CNS tissues were collected at defined disease stages and analyzed for Ninjurin-1 expression by flow cytometry. (**A**) RR-EAE clinical disease course (n = 8). (**B**) CD45^hi^ B220⁻ CD3⁻ CD11b⁺ infiltrating myeloid cells in the CNS displayed a marked increase in Ninjurin-1 expression compared with splenic myeloid cells at all disease stages (pre-onset, onset, peak, remission, and relapse). (**C**) CD45^hi^ CD3⁺ CD11b^-^ T cells in the CNS displayed consistently low Ninjurin-1 expression, with no significant differences compared to splenic CD3⁺ T cells at any disease stage. (**D**) Ninjurin-1 expression in CNS-infiltrating myeloid cells peaked at disease onset and remained elevated at the peak phase. All time points include n = 4 mice, except relapse (n = 3). Data are representative of two independent EAE experiments. Results are shown as mean ± SEM and were analyzed using two-way ANOVA (*p < 0.05, **p < 0.01, ***p < 0.001).

Across all disease phases, Ninjurin-1 expression was significantly higher on infiltrating CNS myeloid cells (CD45^hi^B220⁻CD3⁻CD11b⁺) compared to peripheral myeloid cells in the spleen (**Fig. 1B**). Specifically, Ninjurin-1⁺ myeloid cells accounted for 29% versus 12% at pre-onset, 52% versus 6% at onset, 57% versus 9% at peak, 32% versus 5% at remission, and 36% versus 12% at relapse of the total myeloid cells recorded (CNS vs. spleen, respectively). This enrichment suggests that Ninjurin-1⁺ cells may preferentially migrate to or upregulate Ninjurin-1 expression within the CNS microenvironment.

In contrast, CD3⁺ T cells (CD45^hi^B220⁻CD11b⁻CD3⁺) exhibited consistently low Ninjurin-1 expression throughout RR-EAE, with no significant differences between CNS (3.1 ± 1.5%) and spleen (1.7 ± 1.0%) compartments (**Fig. 1C**).

Together, these results demonstrate that Ninjurin-1 is predominantly expressed by CNS-infiltrating myeloid cells during RR-EAE, with maximal expression observed at disease onset and peak of disease (**Fig. 1D**). These findings identify Ninjurin-1 as a potential marker of inflammatory myeloid cell activation within the CNS during neuroinflammatory progression.

### 3.2 CD4⁺ TH17- and TH1-like conditions increase Ninjurin-1 expression in both BBB endothelial and myeloid cells

Having observed elevated Ninjurin-1 expression on CNS infiltrating myeloid cells, we next investigated the environmental cues that regulate its expression. We previously demonstrated that Ninjurin-1 plays a role in mediating myeloid cell binding to BBB-ECs through homophilic binding [12]. T helper cells, especially TH1 and TH17 subsets, are known to promote inflammation and promote adhesion molecules through their cytokine release in EAE/MS [21, 22, 24], we sought to determine whether cytokines associated these T helper subsets could modulate Ninjurin-1 expression on either BBB-ECs and myeloid cells.

Primary mouse BBB-ECs and splenic CD45⁺CD11b⁺ myeloid cells were incubated for 24 hours under resting conditions or stimulated with individual cytokines: GM-CSF, IFN-γ, IL-1β, IL-4, IL-6, IL-10, IL-17, TGF-β, or TNF-α, or under combined TH1-like (IFN-γ + TNF-α) and TH17-like (IL-17 + GM-CSF) conditions. Ninjurin-1 expression was quantified by flow cytometry relative to resting cells.

On BBB-ECs, only the TH1- and TH17-like conditions significantly increased Ninjurin-1 expression, rising from 13.6% in resting cells to 24.8% and 21.7%, respectively (n = 5; **Fig. 2A**). Similarly, CD11b⁺ myeloid cells isolated from naïve spleens showed marked upregulation of Ninjurin-1 under GM-CSF (14.3%; **p < 0.01), TNF-α (16.1%; ***p < 0.001), TH1-like (13.2%; **p < 0.01), and TH17-like (14.9%; ***p < 0.001) conditions compared to resting cells (8.6%; n = 6; **Fig. 2B**). Other cytokines had no significant effect on Ninjurin-1 expression in either BBB-ECs or myeloid cells.

**Figure 2.**
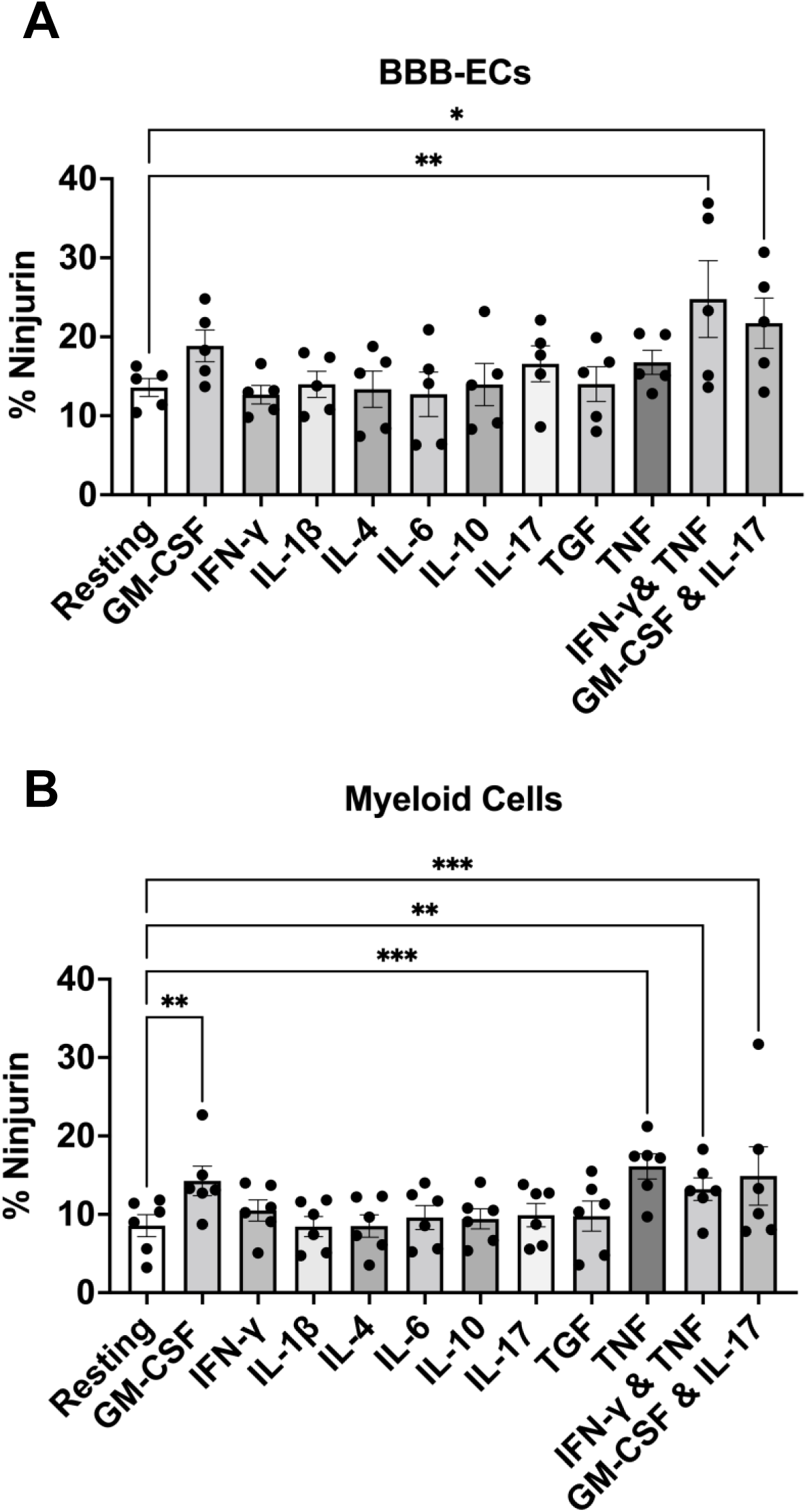
Inflammatory conditions increase Ninjurin-1 expression on BBB endothelial and myeloid cells. Primary brain microvascular endothelial cells (BBB-ECs) and splenic myeloid cells were cultured for 24 hours under resting conditions or in the presence of inflammatory cytokines to assess Ninjurin-1 expression by flow cytometry. (**A**) BBB-ECs upregulated Ninjurin-1 expression following stimulation with TH1-like (IFN-γ + TNF-α) or TH17-like (GM-CSF + IL-17) cytokine combinations (n = 5 independent experiments). (**B**) CD45⁺ CD11b⁺ Ly6G⁻ myeloid cells isolated from spleens of naïve C57BL/6 mice exhibited increased Ninjurin-1 expression when stimulated with GM-CSF, TNF-α, or TH1-/TH17-like cytokine conditions (n = 6 independent experiments). Data are presented as mean ± SEM and analyzed using two-way ANOVA (*p < 0.05, **p < 0.01, ***p < 0.001).

These results align with our previous observations in human BBB-ECs [12], where IFN-γ and TNF-α induced Ninjurin-1 upregulation. Because TH1 and TH17 cytokines are key inflammatory mediators in EAE and MS [24], these findings suggest that cytokine-rich environments characteristic of the inflamed CNS drive Ninjurin-1 upregulation on both endothelial and myeloid cells, potentially facilitating immune cell adhesion and recruitment across the BBB.

### 3.3 Ninjurin-1⁺ myeloid cells exhibit enhanced co-stimulatory and pro-inflammatory profiles in vivo

Given the elevated expression of Ninjurin-1 on CNS infiltrating myeloid cells and their established role in EAE pathology, we next compared the phenotypic and functional characteristics of Ninjurin-1⁺ and Ninjurin-1⁻ myeloid populations. Immune cells were isolated from the CNS and spleen of EAE mice at disease onset (day 11-13) when Ninjurin-1 expression was maximal in the CNS (**Fig. 1D**).

Flow cytometric analysis revealed that CD45^hi^CD11b⁺Ninjurin-1⁺ myeloid cells from the CNS expressed significantly higher levels of the co-stimulatory molecules CD80 and CD86, as well as MHC II, compared with their Ninjurin-1⁻ counterparts (**Fig. 3A**; n = 5). A similar upregulation of these activation markers was also observed in splenic CD11b⁺Ninjurin-1⁺ cells (**Fig. S2A**). PD-L1 expression did not differ significantly between Ninjurin-1⁺ and Ninjurin-1⁻ populations in either compartment.

**Figure 3.**
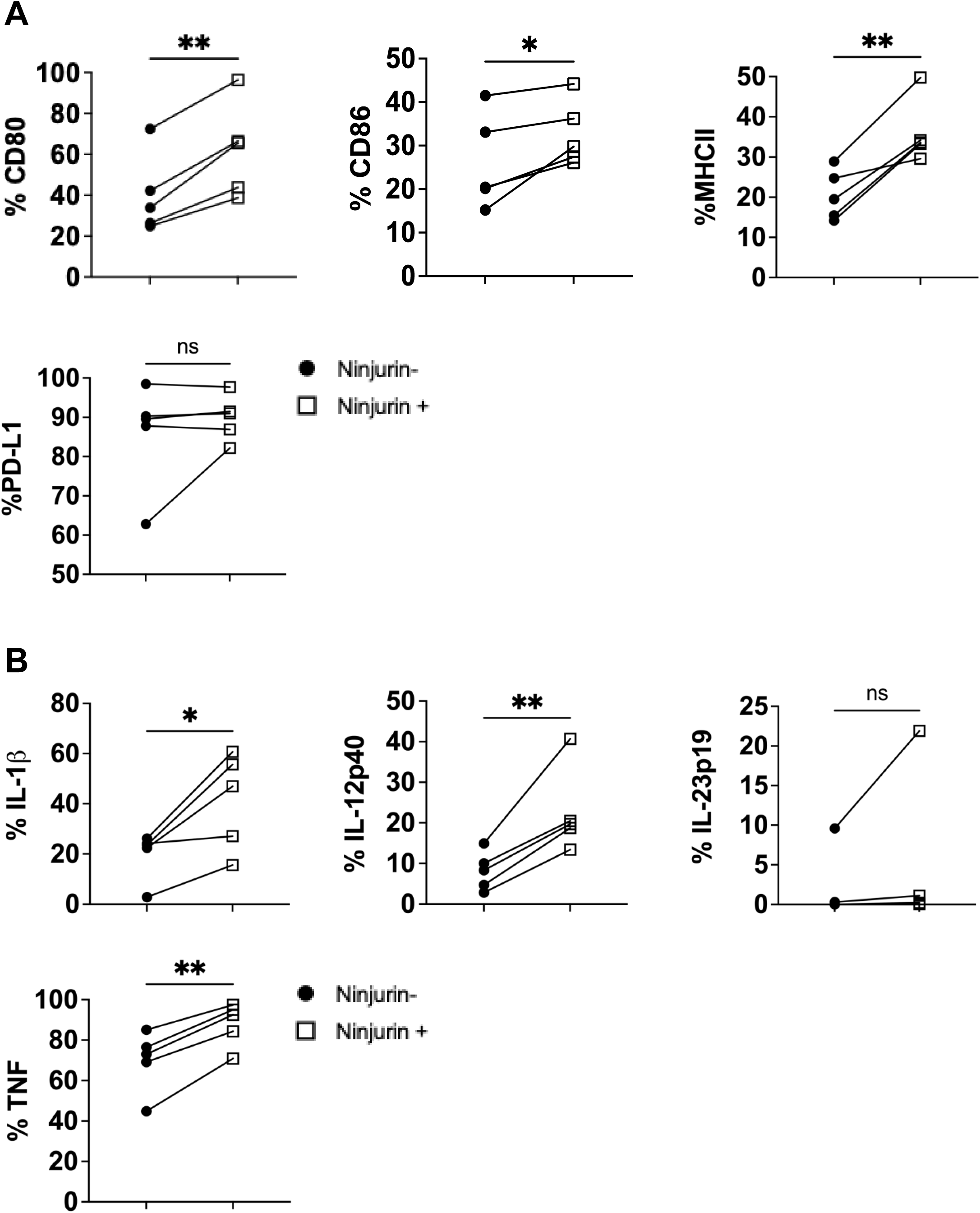
Ninjurin-1⁺ myeloid cells exhibit enhanced activation and inflammatory cytokine expression in the CNS. CD45^hi^ B220⁻ CD11b⁺ infiltrating myeloid cells were isolated from the CNS of SJL/J mice at disease onset during RR-EAE and analyzed by flow cytometry. (**A**) Ninjurin-1⁺ myeloid cells (open squares) displayed significantly higher expression of the co-stimulatory molecules CD80 and CD86, as well as MHC class II, compared with Ninjurin-1⁻ counterparts (closed circles). (**B**) Ninjurin-1⁺ myeloid cells also exhibited increased expression of the inflammatory cytokines TNF-α, IL-1β, and IL-12p40. Data represent n = 5 mice and are presented as mean ± SEM. Statistical significance was determined using paired t-test (*p < 0.05, **p < 0.01).

We next assessed the expression of inflammatory cytokines associated with TH1 and TH17 polarization, including TNF-α, IL-1β, IL-12p40, and IL-23p19. In the CNS, Ninjurin-1⁺ myeloid cells expressed significantly higher levels of TNF-α, IL-1β, and IL-12p40 than Ninjurin-1⁻ cells (**Fig. 3B**). In the spleen, Ninjurin-1⁺ myeloid cells also exhibited increased expression of IL-1β, IL-6, IL-12p40, IL-23p19, TGF-β, and TNF-α (**Fig. S2B**).

Together, these findings indicate that Ninjurin-1⁺ myeloid cells display a heightened activation state characterized by elevated co-stimulatory molecule expression and robust production of pro-inflammatory cytokines. This phenotype suggests an enhanced capacity of Ninjurin-1⁺ cells to promote T-cell activation and sustain CNS inflammation during RR-EAE.

### 3.4 Ninjurin-1⁺ CD11b⁺Ly6G⁻ splenocytes exhibit differential expression of antigen-presentation and inflammatory genes

Given the marked differences in co-stimulatory molecule and cytokine expression between Ninjurin-1⁺ and Ninjurin-1⁻ myeloid cells, we next investigated whether these populations possess distinct transcriptional profiles independent of disease context. To this end, CD45⁺CD11b⁺B220⁻CD3⁻Ly6G⁻Ninjurin-1⁺ and CD45⁺CD11b⁺B220⁻CD3⁻Ly6G⁻Ninjurin-1⁻ splenocytes were flow-sorted from naïve mice (gating shown in **Fig. S3**). RNA was extracted, converted to cDNA, and subjected to targeted qPCR profiling of genes involved in antigen presentation and innate immune activation.

Overall, the transcriptional data support the hypothesis that Ninjurin-1⁺ myeloid cells are intrinsically more pro-inflammatory than their Ninjurin-1⁻ counterparts. Genes associated with antigen presentation and immune activation, including Tnf, Cd40, H2-DMa, and Relb, were significantly upregulated in Ninjurin-1⁺ cells (**Table 1**). Enhanced expression of Tlr1 and Tlr7 further indicates increased responsiveness to innate immune stimuli. In parallel, elevated expression of adhesion and migration molecules such as Icam1 and Cd44 suggests that Ninjurin-1⁺ cells are well equipped to traffic to and interact with inflamed tissues, reinforcing their potential role in amplifying CNS inflammation.

**Table 1.**
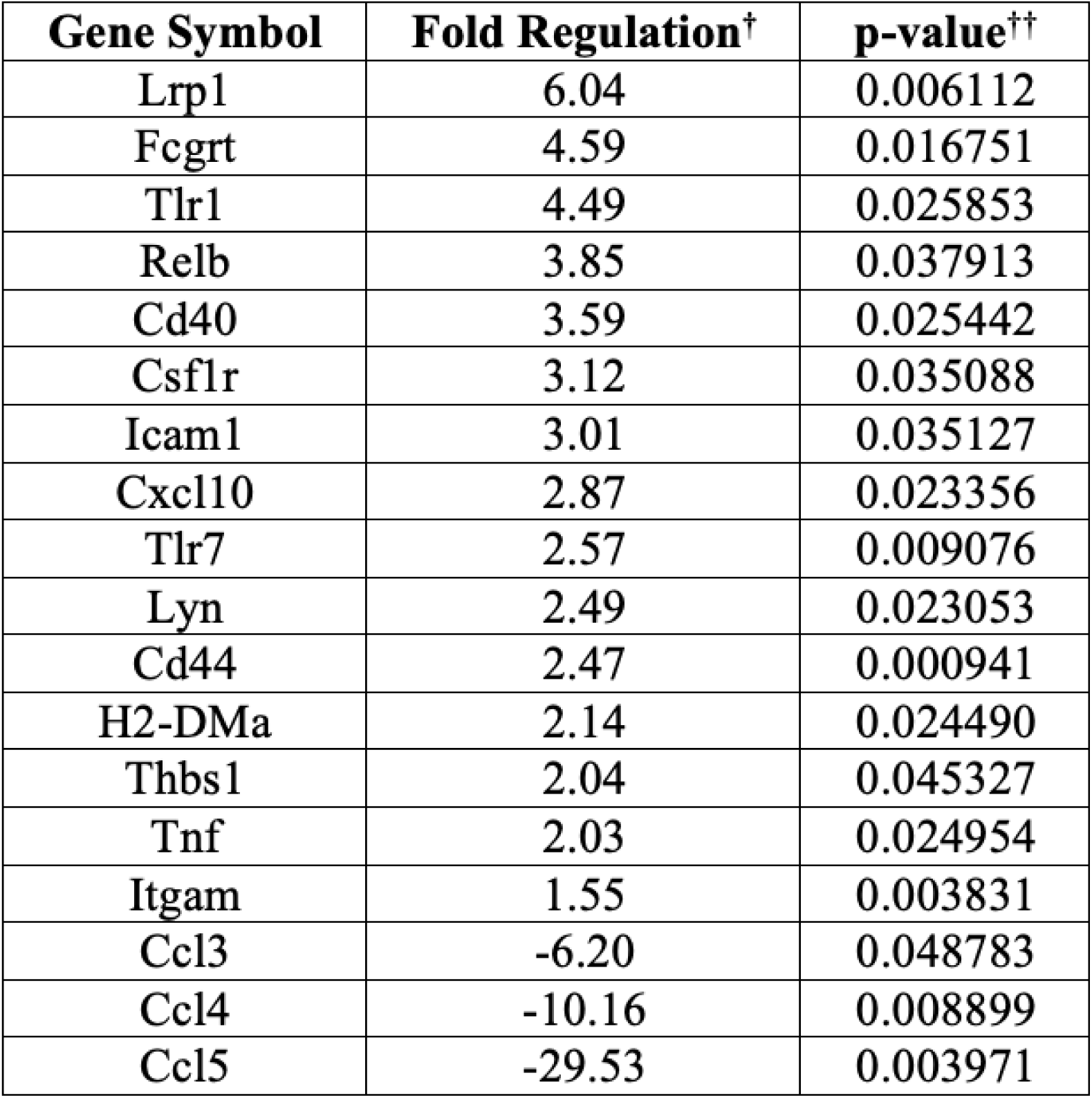
List of genes differentially expressed between Ninjurin-1⁺ and Ninjurin-1^-^myeloid cells. Shown are genes with their corresponding fold-regulation values and *p*-values comparing Ninjurin-1⁺ versus Ninjurin-1^-^ CD45⁺CD11b⁺B220^-^CD3^-^Ly6G^-^ myeloid cells, as determined by RT² Profiler PCR Array analysis. ^†^ Only genes showing ≥1.5-fold difference between groups are represented. Positive values indicate upregulation in Ninjurin-1⁺ cells relative to Ninjurin-1^-^ cells. ^††^ *p*-values were calculated using a Student’s *t*-test on replicate 2^(−ΔCT) values for each gene in the Ninjurin-1^+^ and Ninjurin-1^-^ groups (*n* = 3 independent experiments).

Interestingly, several chemokines involved in leukocyte recruitment, including Ccl3, Ccl4, and Ccl5 (RANTES), were downregulated, whereas Cxcl10 (IP-10) was upregulated, indicating a shift toward a more selective chemokine profile that may favor specific immune-cell interactions within inflammatory microenvironments.

Collectively, these transcriptional data indicate that Ninjurin-1 expression defines a subset of myeloid cells with heightened antigen-presenting capacity, migratory potential, and pro-inflammatory gene expression, positioning Ninjurin-1 as both a functional adhesion molecule and a marker of inflammatory myeloid activation.

### 3.5 Ninjurin-1⁺ myeloid cells enhance activation and proliferation of MOG_35-55_-specific CD4⁺ T cells

Building on our previous findings, we hypothesized that Ninjurin-1⁺ myeloid cells act as more potent antigen-presenting cells (APCs), leading to stronger CD4⁺ T cell activation. To test this, we co-cultured FACS-sorted Ninjurin-1⁺ or Ninjurin-1⁻ myeloid cells with MOG_35-55_ peptide (20 μg/ml) and CFSE-labeled CD4⁺ T cell isolated from 2D2 transgenic mice, which express a T cell receptor specific for MOG_35-55_. 2D2 T cells were also cultured with MOG_35-55_ with no myeloid cells and co-cultured with myeloid cells without MOG_35-55_ as controls. After 72 hours of co-culture, T cell proliferation and phenotype was assessed by CFSE dilution and expression of CD25, IL-17, GM-CSF, IFN-γ, and TNF-α.

CD4⁺ T cells co-cultured with Ninjurin-1⁺ APCs displayed markedly greater proliferation compared to those cultured with Ninjurin-1⁻ cells (**Fig. 4A**). This enhanced proliferation was accompanied by increased CD25 expression, consistent with elevated T cell activation (**Fig. 4B**). Furthermore, cytokine profiling revealed that CD4⁺ T cells stimulated by Ninjurin-1⁺ APCs produced higher levels of TNF-α (**Fig. 4B**), suggesting a stronger pro-inflammatory response.

**Figure 4.**
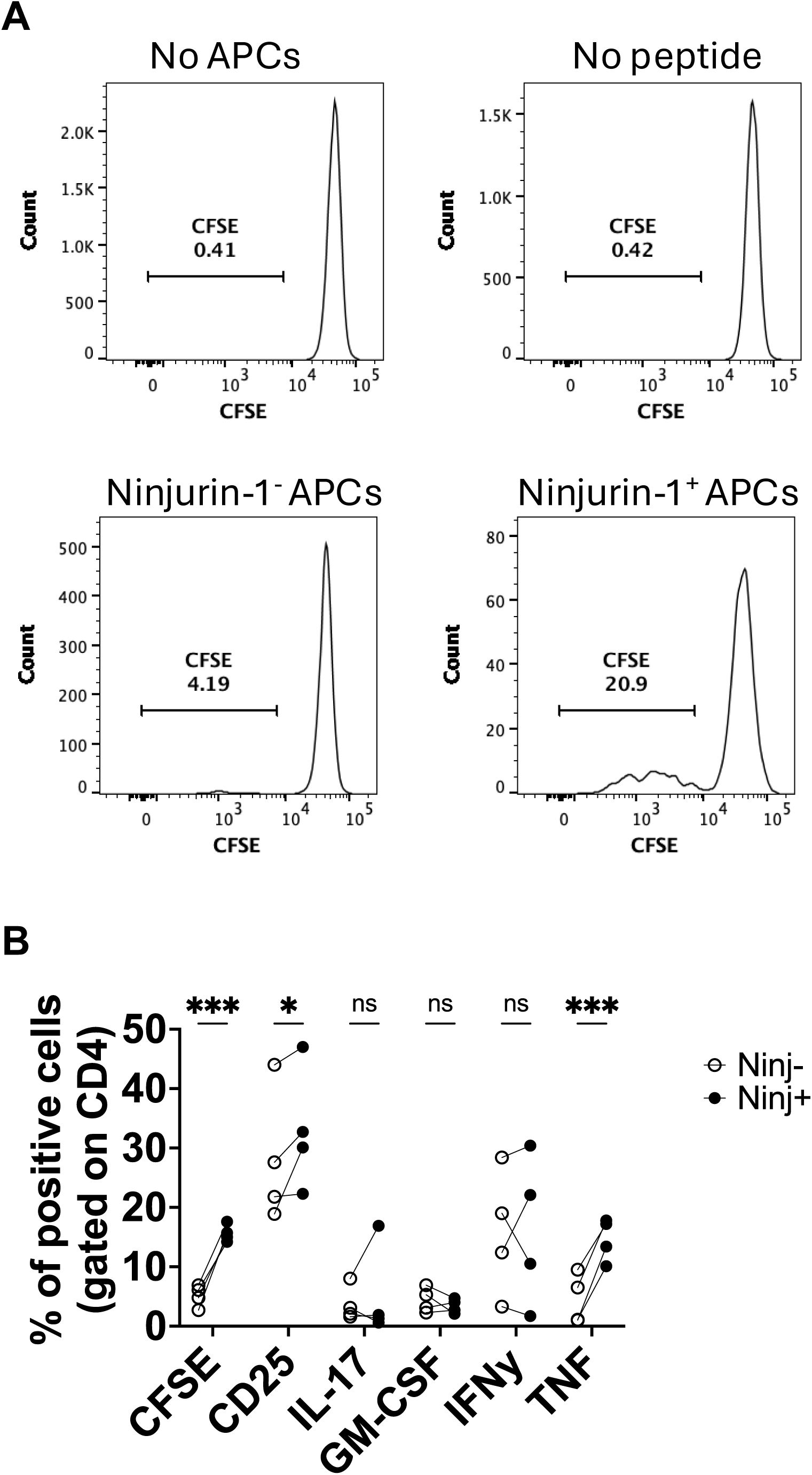
Ninjurin-1⁺ myeloid cells enhance CD4⁺ T cell activation and cytokine production. CD45⁺ B220⁻ CD3⁻ CD11b⁺ Ly6G⁻ Ninjurin-1⁺ or Ninjurin-1⁻ myeloid cells were flow-sorted and co-cultured with CFSE-labeled CD4⁺ T cells isolated from 2D2 mice. Cells were stimulated with MOG_35-55_ peptide in 96-well plates for 72 hours. (**A**) T cell proliferation, measured by CFSE dilution, was greater when CD4⁺ T cells were co-cultured with Ninjurin-1⁺ myeloid cells compared to Ninjurin-1⁻ cells. Controls included CD4⁺ T cells cultured without antigen-presenting cells (APCs) or without MOG_35-55_ peptide. (**B**) CD4⁺ T cells co-cultured with Ninjurin-1⁺ myeloid cells showed increased expression of CD25 and TNF-α, indicating higher levels of activation and inflammatory cytokine production. Data represent n = 4 independent experiments and are shown as mean ± SEM. Statistical analysis was performed using two-way ANOVA (*p < 0.05, **p ≤ 0.01, ***p < 0.001).

Together, these results demonstrate that Ninjurin-1⁺ myeloid cells exhibit superior antigen-presenting capacity and promote a more robust and inflammatory CD4⁺ T cell response, further supporting their role as key drivers of neuroinflammation during EAE.

### 3.6 Anti-Ninjurin-1 treatment at peak disease ameliorates RR-EAE

Given that Ninjurin-1 remains highly expressed in the CNS throughout all phases of RR-EAE, we next tested whether blocking its function could modify disease progression. To this end, SJL/J mice were treated intraperitoneally with 1 mg/mL of the anti-Ninjurin-1_26-37_ blocking peptide or a scramble control three times per week, beginning at the peak of disease (∼day 15). Mice were monitored and disease severity was scored daily until day 30, at which point the CNS was harvested for histological and flow cytometric analyses.

Mice treated with anti-Ninjurin-1 displayed significantly lower clinical scores and were largely protected from relapse, indicating a marked improvement in disease outcome (**Fig. 5A**). Consistent with the known role of Ninjurin-1 as an adhesion molecule, its blockade resulted in a substantial reduction in immune cell infiltration into the CNS, including total CD45⁺ leukocytes, CD3⁺ and CD4⁺ T cells, B220⁺ B cells, and CD11b⁺ myeloid cells (**Fig. 5B**).

**Figure 5.**
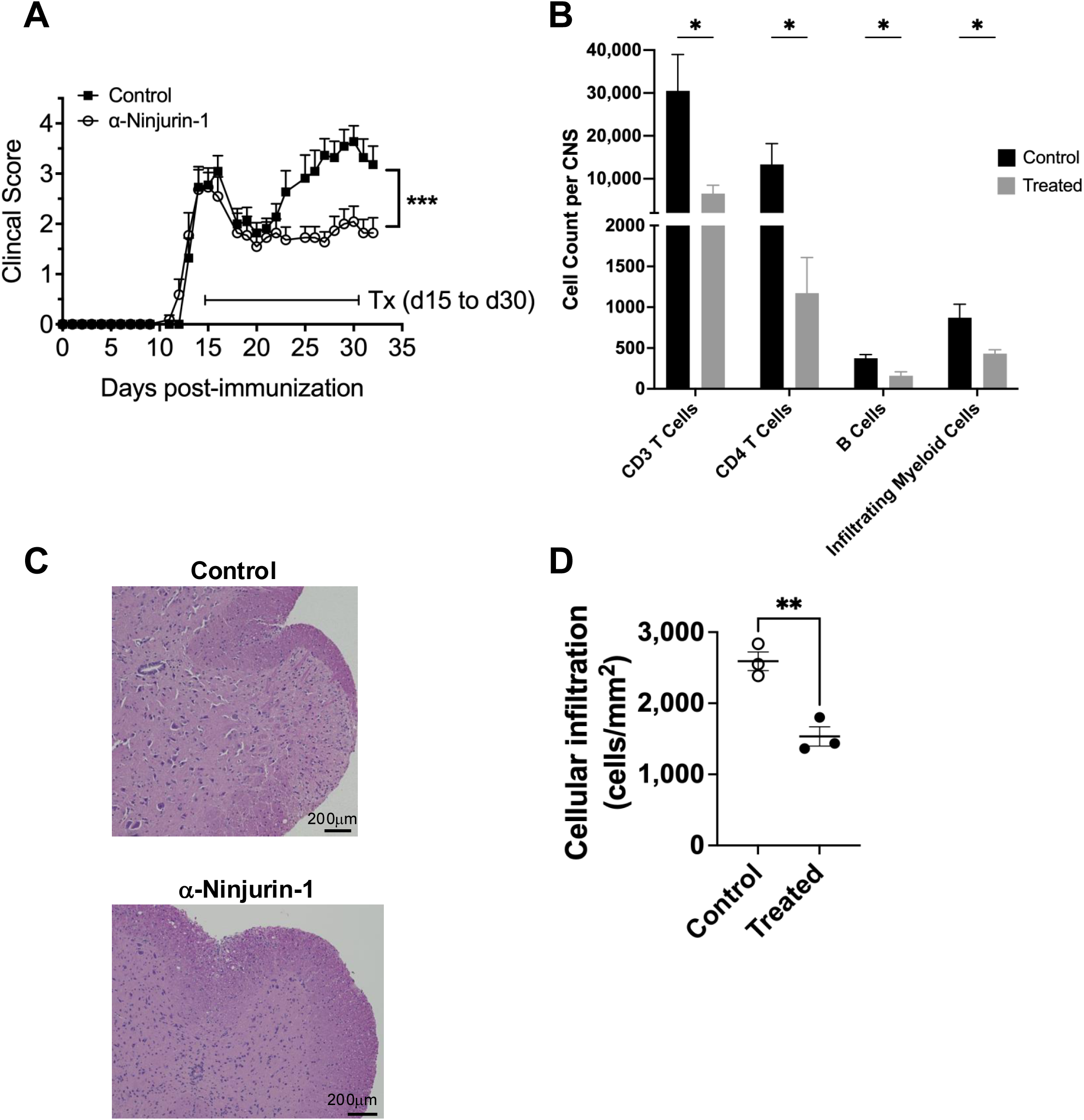
Anti-Ninj_26–37_ treatment at peak RR-EAE protects against relapse and reduces CNS immune cell infiltration. RR-EAE was induced in SJL/J mice by subcutaneous injection of PLP_139-151_ emulsified in CFA. Mice were treated intraperitoneally three times per week with anti-Ninj_26–37_ peptide (1 mg/mL; n = 11) or scramble control (n = 11), starting at the peak of disease (day 15) and continuing until day 30 post-induction. (**A**) Anti-Ninj_26–37_–treated mice (open circles) displayed significantly reduced clinical scores and were protected from relapse compared with scramble-treated controls (closed squares; ***p < 0.001 by Mann–Whitney test). (**B**) Flow cytometry of CNS mononuclear cells showed a marked reduction in total CD45^hi^ leukocytes, including CD3⁺ T cells, CD4⁺ T cells, B220⁺ B cells, and CD11b⁺ myeloid cells, in anti-Ninj_26–37_–treated mice (n = 5; *p < 0.05 by multiple unpaired t-tests). (**C**) Representative H&E-stained spinal cord sections collected at day 30 show reduced inflammatory infiltrates in anti-Ninj_26–37_–treated mice compared to controls. Data are representative of at least three sections per mouse from four mice per group. Scale bar = 200 µm. (**D**) Cellular infiltration was quantified by counting hematoxylin-positive nuclei within standardized 300 × 300 µm white matter regions (three ROIs per section, averaged per mouse) and expressed as cells per mm². Each dot represents one mouse; bars indicate mean ± SEM.

Histological assessment at the end of treatment confirmed these findings, as spinal cord sections from anti-Ninjurin-1-treated mice exhibited fewer immune cell infiltrates compared to controls (**Fig. 5C**). Quantification of cellular infiltration in H&E-stained spinal cord sections revealed a significant reduction in anti-Ninjurin-1-treated EAE mice compared with controls, as assessed by nuclear density within standardized white matter regions (**Fig. 5D**; cells/mm²; n = 3 mice per group). These results are consistent with previous studies reporting reduced APC infiltration in chronic EAE following Ninjurin-1 blockade, supporting a role for Ninjurin-1 in mediating immune cell adhesion and trafficking into the CNS.

Together, these findings demonstrate that therapeutic blockade of Ninjurin-1 ameliorates RR-EAE when administered at peak disease, at least in part by limiting CNS immune infiltration and inflammation, thereby supporting Ninjurin-1 as a promising immunomodulatory target for relapsing-remitting MS.

## 4. Discussion

This study identifies Ninjurin-1 as a critical regulator of neuroinflammation in RR-EAE, revealing its dual roles as both an adhesion molecule and a marker of inflammatory activation in myeloid cells. We show that Ninjurin-1 expression is markedly upregulated in the CNS during active disease and that its blockade mitigates immune infiltration and clinical severity. These findings establish Ninjurin-1 as a previously unrecognized mediator linking peripheral immune activation to central neuroinflammation.

Our data demonstrates that Ninjurin-1 expression defines a population of myeloid cells with enhanced pro-inflammatory and antigen-presenting functions. Ninjurin-1⁺ myeloid cells exhibited higher expression of MHC II and co-stimulatory molecules, increased production of cytokines such as TNF-α and IL-1β, and a distinct transcriptional profile enriched for genes associated with antigen presentation, Toll-like receptor signaling, and cell adhesion. The upregulation of Tlr1 and Tlr7 in these cells is particularly relevant, as both receptors have been implicated in MS and EAE [25, 26]. Activation of TLR1 enhances microglial and macrophage responsiveness to bacterial lipoproteins and misfolded proteins, and promotes pro-inflammatory cytokine release [27, 28], while TLR7 stimulation induces type I interferons and TNF-α, contributing to CNS inflammation and demyelination [29, 30]. Thus, Ninjurin-1⁺ cells may represent a population highly responsive to TLR-driven inflammatory cues within the CNS, further amplifying innate and adaptive immune activation. Together, these transcriptional changes outline a multifaceted activation program in Ninjurin-1⁺ myeloid cells that integrates innate sensing, adhesion, and cytokine signaling.

The concurrent upregulation of Icam1 and Cd44 suggests that these cells are primed for tissue infiltration and sustain interaction with other immune cells within the inflamed CNS. Interestingly, Cxcl10 (IP-10), a chemokine known to recruit CXCR3⁺ Th1 cells across the BBB [31], was also increased in Ninjurin-1⁺ myeloid cells. Elevated CXCL10 levels have been observed in active MS lesions and in EAE, where they promote the accumulation of pathogenic T cells and perpetuate the inflammatory milieu [32, 33]. These features parallel pathogenic myeloid subsets identified in MS lesions, which have been shown to drive demyelination and neurodegeneration [34–37], and support a model in which Ninjurin-1⁺ myeloid cells drive lesion formation and disease progression through both antigen presentation and chemokine-mediated immune recruitment.

Importantly, we found that TH1- and TH17-associated cytokines, including IFN-γ, TNF-α, GM-CSF, and IL-17 induce Ninjurin-1 expression on both BBB-ECs and myeloid cells. This places Ninjurin-1 downstream of key inflammatory pathways known to dominate during MS relapses. Such cytokine-driven upregulation provides a mechanistic link between systemic immune activation and localized BBB disruption, where Ninjurin-1 may facilitate leukocyte adhesion and transmigration. The observed upregulation of Ninjurin-1 on both endothelial and infiltrating myeloid cells supports the concept of homophilic interactions across the BBB, potentially stabilizing immune cell adhesion and promoting CNS entry.

Therapeutic blockade of Ninjurin-1 at the peak of disease attenuated clinical scores, reduced immune cell infiltration, and decreased CNS inflammation. This effect resembles the efficacy of adhesion-molecule inhibitors such as Natalizumab, which targets α4-integrin/VCAM-1 interactions [38]. However, Ninjurin-1’s unique dual role distinguishes it from canonical adhesion molecules. Unlike ICAM-1 and VCAM-1, which primarily mediate leukocyte-endothelial interactions, Ninjurin-1 exerts a dual influence by functioning both on vascular and immune compartments. Its homophilic binding supports leukocyte adhesion at the BBB, while its intracellular signaling capacity enhances myeloid activation and antigen presentation within the CNS. This combination positions Ninjurin-1 as both a structural and signaling mediator of neuroinflammation, extending its role beyond passive adhesion to active immune modulation.

Beyond its adhesive role, Ninjurin-1 may also influence inflammatory cell survival and signaling. Previous reports have implicated Ninjurin-1 in plasma membrane rupture and the subsequent release of damage-associated molecular patterns [16, 39, 40], processes that can propagate sterile inflammation. Together with our findings, this raises the possibility that Ninjurin-1 contributes to a feed-forward loop wherein inflammatory cytokines induce its expression, promoting adhesion, activation, and additional cytokine release that sustain neuroinflammation.

Our findings extend the understanding of adhesion molecules in MS by emphasizing immune cell-intrinsic regulation rather than endothelial expression alone. By integrating cytokine signaling, adhesion, and immune activation, Ninjurin-1 bridges molecular pathways that have traditionally been studied in isolation. Given that Ninjurin-1 is upregulated in multiple inflammatory conditions, including stroke and fibrosis, its role may not be limited to demyelinating disease but may represent a broader mechanism of leukocyte-mediated tissue injury.

In conclusion, we identify Ninjurin-1 as a novel adhesion and activation molecule driving myeloid-mediated inflammation in RR-EAE. Through its induction by TH1/TH17 cytokines and expression on both BBB-endothelial and immune cells, Ninjurin-1 orchestrates key steps in immune cell recruitment and activation within the CNS. Its blockade alleviates disease even at advanced stages, underscoring its therapeutic potential. Future studies should assess Ninjurin-1 expression in human MS lesions and determine whether its signaling partners and downstream pathways can be targeted to modulate disease progression. Elucidating the intracellular signaling cascades engaged by Ninjurin-1 will be essential to understand how this molecule bridges adhesion and immune activation, and may uncover new opportunities for therapeutic intervention.

## Author Contributions

C.T. conducted most the experiments. A.Annett assisted with EAE studies. C.T., A. Alkhimovitch, and K.S.P had intellectual input and revised the manuscript. I.I. supervised the project and revised the manuscript. All authors contributed to manuscript preparation.

## Acknowledgments

I.I. is supported by the National Institutes of Health (NIH) through an R35 Maximizing Investigators’ Research Award (R35GM146890) and an R03 grant (R03NS144870). We thank the University of Cincinnati’s histology core and the Laboratory Animal Medical Services (LAMS), as well as Cincinnati Children’s Hospital flow cytometry core for technical support.

