## Supplemental Information for "Dual Role of Ninjurin-1 in Myeloid Cell Adhesion and Inflammation in Relapse-Remitting EAE"

#### FIGURE LEGENDS

**Figure S1. Gating strategy for CNS immune cells and Ninjurin-1 detection using isotype control.** (**A**) CNS single-cell suspensions were first gated by forward and side scatter (FSC/SSC; P1), followed by exclusion of doublets and dead cells. Infiltrating myeloid cells were identified as $CD45^{hi}$ $CD11b^{+}$ and infiltrating lymphoid cells as $CD45^{hi}$ $CD11b^{-}$. Within the lymphoid compartment, $CD3^{+}$ T cells were further defined. (**B**) Ninjurin-1 expression was assessed on infiltrating myeloid cells and $CD3^{+}$ T cells. Representative plots show isotype control staining (black) and Ninjurin-1 staining (red).

**Figure S2. Increased activation and cytokine expression by Ninjurin-1$^{+}$ myeloid cells in the spleen.** Comparison of Ninjurin-1$^{-}$ (closed circles) and Ninjurin-1$^{+}$ (open squares) $CD45^{hi}$ $B220^{-}$ $CD11b^{+}$ myeloid cells isolated from the spleen at onset of RR-EAE. Flow cytometry analysis revealed that (**A**) co-stimulatory molecules CD80 and CD86, as well as MHC II, were significantly increased on Ninjurin-1$^{+}$ cells. (**B**) Expression of IL-1β, IL-6, IL-12p40, IL-23p19, TGF-β, and TNF-α was also elevated in Ninjurin-1$^{+}$ myeloid cells ($n = 6$ mice; $^{*}p < 0.05$, $^{**}p < 0.01$, $^{***}p < 0.001$ by paired t-test).

**Figure S3. Cell sorting of $CD45^{+}CD11b^{+}B220^{-}CD3^{-}Ly6G^{-}$Ninjurin-1$^{+}$ and Ninjurin-1$^{-}$ myeloid cells.** (**A**) Splenocytes were first enriched for $CD11b^{+}$ cells using Miltenyi magnetic microbeads, followed by sorting on a Sony MA900 cell sorter. At the FACS, cells were sequentially gated by forward and side scatter (FSC/SSC), single-cell discrimination, and viability. Myeloid cells were identified as $CD45^{+}CD11b^{+}CD3^{-}B220^{-}$, with $Ly6G^{+}$ neutrophils excluded. (**B**) Ninjurin-1$^{+}$ and Ninjurin-1$^{-}$ populations were defined using an isotype control to establish gating thresholds.

### Figure S1

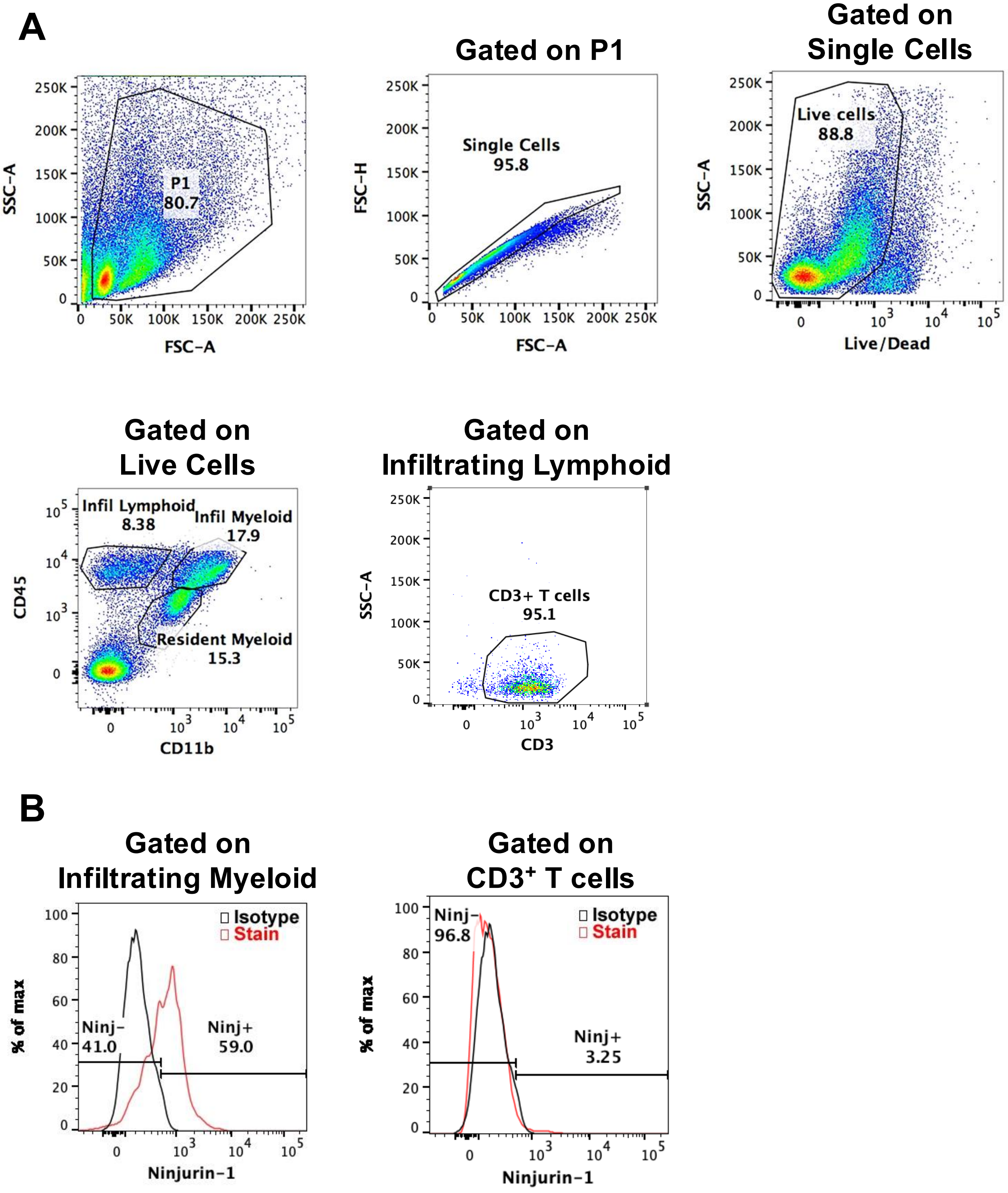
A
Gated on P1
Gated on Single Cells
P1
80.7
SSC-A
FSC-A
Single Cells
95.8
FSC-H
Live cells
88.8
Live/Dead
Gated on Live Cells
Gated on Infiltrating Lymphoid
Infil Lymphoid
8.38
Infil Myeloid
17.9
Resident Myeloid
15.3
CD45
CD11b
CD3+ T cells
95.1
CD3
B
Gated on Infiltrating Myeloid
Gated on CD3+ T cells
Isotype
Stain
Ninj-
41.0
Ninj+
59.0
% of max
Ninjurin-1
Ninj-
96.8
Ninj+
3.25

### Figure S2

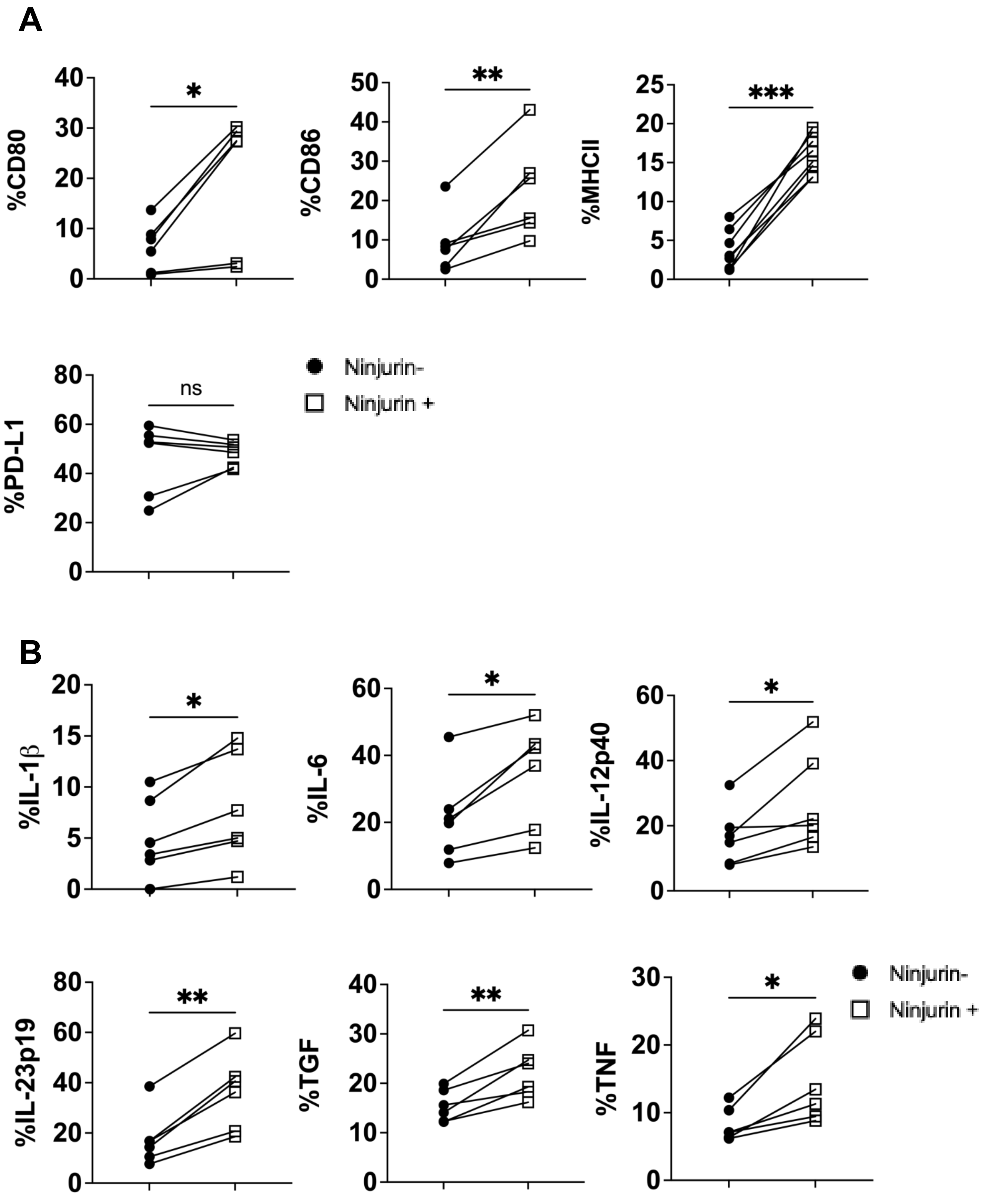

A
%CD80
%CD86
%MHCII
%PD-L1
*
**
***
ns
Ninjurin-
Ninjurin +
B
%IL-1β
%IL-6
%IL-12p40
%IL-23p19
%TGF
%TNF
*
**
Ninjurin-
Ninjurin +

### Figure S3

**A**

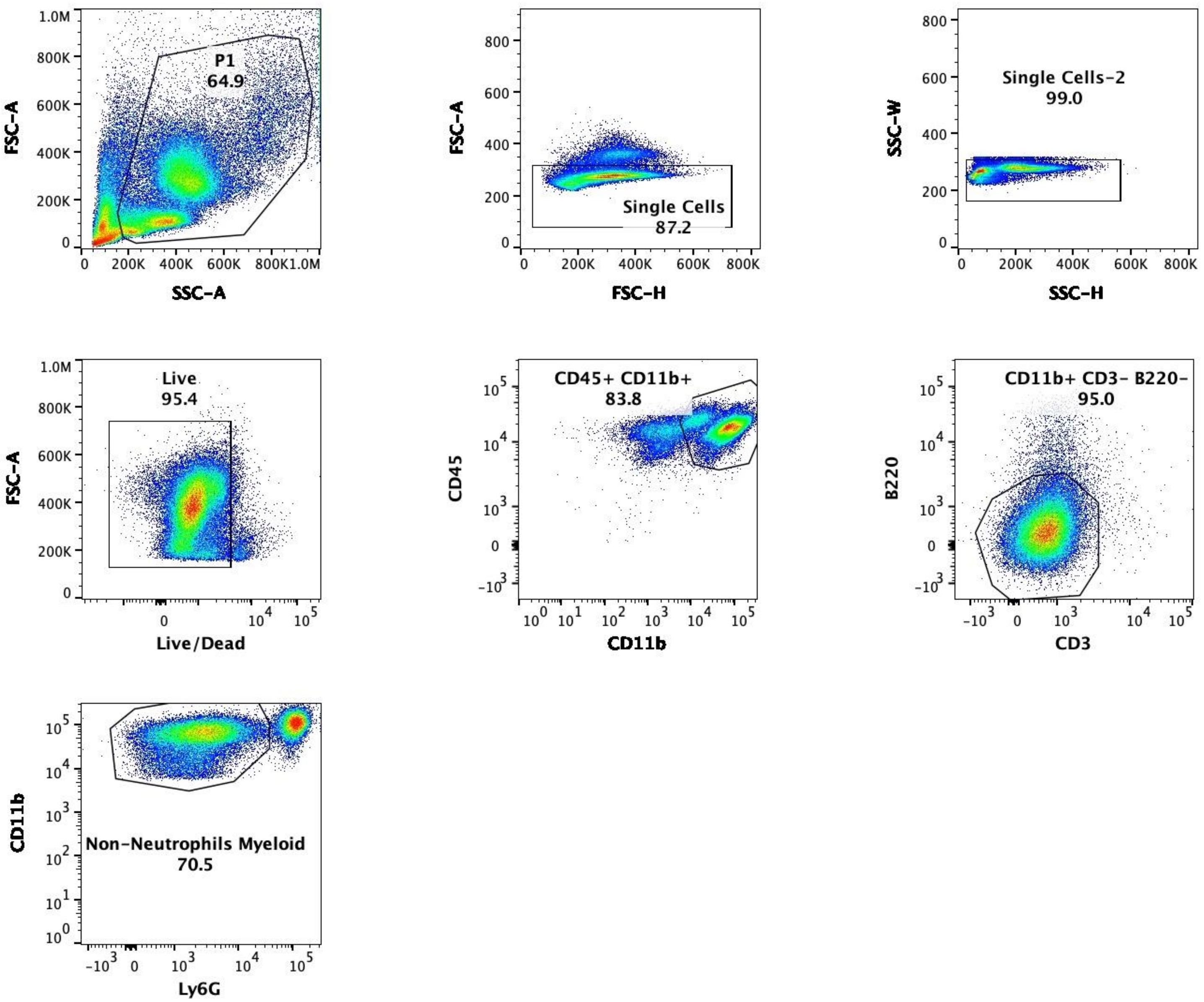
P1
64.9
FSC-A
SSC-A
Single Cells
87.2
FSC-H
Single Cells-2
99.0
SSC-W
SSC-H
Live
95.4
Live/Dead
CD45+ CD11b+
83.8
CD45
CD11b
CD11b+ CD3- B220-
95.0
B220
CD3
Non-Neutrophils Myeloid
70.5
Ly6G

**B**

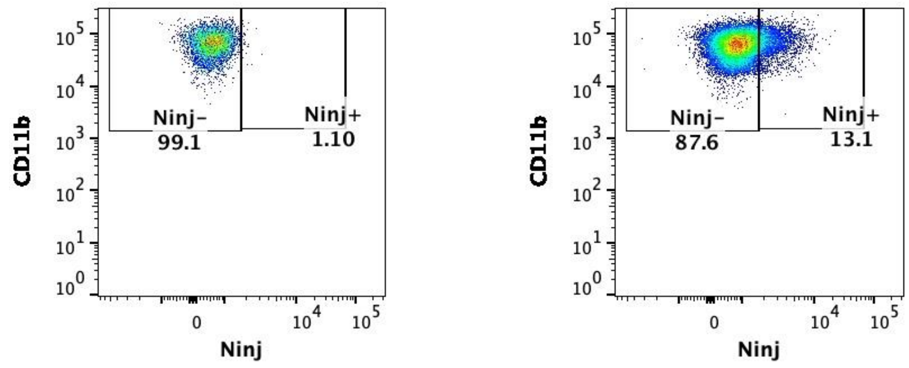
Ninj-
99.1
Ninj+
1.10
Ninj-
87.6
Ninj+
13.1
CD11b
Ninj
